# Recurrent loss of heterozygosity correlates with clinical outcome in pancreatic neuroendocrine cancer

**DOI:** 10.1101/214585

**Authors:** Ben Lawrence, Cherie Blenkiron, Kate Parker, Peter Tsai, Sandra Fitzgerald, Paula Shields, Tamsin Robb, Mee Ling Yeong, Nicole Kramer, Sarah James, Mik Black, Vicky Fan, Nooriyah Poonawala, Patrick Yap, Esther Coats, Braden Woodhouse, Reena Ramsaroop, Masato Yozu, Bridget Robinson, Kimiora Henare, Jonathan Koea, Peter Johnston, Richard Carroll, Saxon Connor, Helen Morrin, Marianne Elston, Christopher Jackson, Papaarangi Reid, John Windsor, Andrew MacCormick, Richard Babor, Adam Bartlett, Dragan Damianovich, Nicholas Knowlton, Sean Grimmond, Michael Findlay, Cristin Print

**Affiliations:** Discipline of Oncology, Faculty of Medicine and Health Sciences, University of Auckland, New Zealand; Department of Molecular Medicine and Pathology, School of Medical Sciences, Faculty of Medicine and Health Sciences, and Maurice Wilkins Centre, University of Auckland, New Zealand; Anatomic Pathology Services, Auckland, New Zealand; LabPlus, Auckland City Hospital, New Zealand; Department of Biochemistry, University of Otago, Dunedin, NZ; Bioinformatics Institute, University of Auckland, New Zealand; Genetic Health Service New Zealand (Northern Hub), Auckland, NZ; Waitemata District Health Board, Auckland, New Zealand; Histopathology Department, Middlemore Hospital, Auckland, New Zealand; Canterbury District Health Board, Christchurch, New Zealand; Auckland Cancer Society Research Centre, Faculty of Medical and Health Sciences, The University of Auckland, New Zealand; Upper Gastrointestinal Unit, Department of Surgery, North Shore Hospital, Takapuna, New Zealand; Department of Surgery, Auckland District Health Board, Auckland, New Zealand; Endocrine, Diabetes and Research Centre, Wellington Regional Hospital, Wellington, New Zealand; Department of Pathology, University of Otago Christchurch, New Zealand; Waikato Clinical Campus, University of Auckland Department of Medicine, New Zealand; Department of Medicine, Dunedin School of Medicine, University of Otago, New Zealand; Te Kupenga Hauora Maori, Faculty of Medical and Health Sciences, University of Auckland, New Zealand; Department of Surgery, University of Auckland, New Zealand; Department of Surgery, Counties Manukau District Health Board, Auckland, New Zealand; Department of Medical Oncology, Auckland City Hospital, Auckland, New Zealand; University of Melbourne Centre for Cancer Research, University of Melbourne, Melbourne, Victoria, Australia

**Keywords:** neuroendocrine cancer, pancreatic, aneuploidy, genomic, pathology

## Abstract

Pancreatic neuroendocrine tumors (pNETs) are uncommon cancers arising from pancreatic islet cells. Analysis of gene mutation, copy number and RNA expression of 57 sporadic pNETs showed that pNET genomes are dominated by aneuploidy. Remarkably, ~25% of pNETs had genomes characterized by recurrent loss of heterozygosity (LoH) of the same 10 chromosomes, accompanied by bi-allelic *MEN1* inactivation, and these cases had generally poor clinical outcome. Another ~25% of all pNETs had chromosome 11 LoH and bi-allelic *MEN1* inactivation, lacking the recurrent LoH pattern – these had universally good clinical outcome. Some level of aneuploidy was common, and overall ~80% of pNETs had LoH of ≥1 chromosome. This aneuploidy led to changes in RNA expression at the level of whole chromosomes and allowed pathogenic germline variants (e.g. *ATM*) to be expressed unopposed, inactivating downstream tumor suppressor pathways. Some pNETs appear to utilize *VHL* gene methylation or mutation to activate pseudo-hypoxia. Contrary to expectation neither tumor morphology within well-differentiated pNETs nor single gene mutation had significant associations with clinical outcome, nor did expression of RNAs reflecting the activity of immune, differentiation, proliferative or tumor suppressor pathways. *MEN1* was the only statistically significant recurrently mutated driver gene in pNETs. Only one pNET had clearly oncogenic and actionable SNVs (in *PTEN* and *FLCN*) confirmed by corroborating RNA expression changes. The two distinct patterns of aneuploidy described here, associated with markedly poor and good clinical outcome respectively, define a novel oncogenic mechanism and the first route to genomic precision oncology for this tumor type.

## Introduction

Pancreatic neuroendocrine tumors (pNETs) are clinically heterogeneous tumors derived from neuroendocrine cells of pancreatic islets, which differ from one other by their primary organ of origin and degree of cellular differentiation. Currently, therapeutic decisions must be made with little knowledge of the biological drivers of individual NETs, underlining the importance of improved genomic understanding of these tumors. Although driver mutations in tumor suppressor genes have been found in pNETs (e.g., *MEN1*, *DAXX*, *ATRX*, *VHL*, *YY1*, and mechanistic target of rapamycin (mTOR) pathway genes (1–3)) they are infrequent and are not generally able to indicate specific systemic therapies. In addition to these driver mutations, other genomic changes have been observed in pNETs, including: telomeric dysregulation (4, 5), copy number (CN) changes (6), changes in RNA expression that indicate mTOR pathway activation (7), germline *MEN1* and *MUTYH* inactivation (2, 8). Epigenetic changes in methylation (9) and microRNA expression (10) have been described in pNETs, with insulinomas especially enriched for changes to the sequence, methylation and expression of genes encoding epigenetic modifiers (11).

Therefore, we undertook pathological examination and deep genomic analysis of a group of clinically homogenous sporadic pNETs. Our results show that pNETs are dominated by aneuploidy along with *MEN1* gene mutation, with mutations evident in small numbers of other genes associated with chromosomal stability. By combining multi-modal genomic information, we found that the extensive loss of heterozygosity (LoH) seen in pNETs is linked to dysregulation of RNA expression on the scale of whole chromosomes, which appears to have driven tumor development with few other genome lesions. Neither patterns of gene mutation or morphology, nor RNA expression patterns reflecting immune, differentiation, proliferative or tumor suppressor pathways had significant association with clinical outcome. However, distinct patterns of recurrent chromosome-level aneuploidy with concordant chromosome-level gene expression changes identified two distinct tumor clusters, which have markedly good and poor clinical outcome. Although the mechanisms that underpin this aneuploidy remain unclear, these two tumor clusters incorporate half of pNETs and will inform clinical care.

## Results

We analyzed 57 sporadic pNETs collected from 53 individuals along with matched normal tissues. Key population characteristics of the pNET series are described in Supplementary Table S1, with individual patient characteristics described in Supplementary Table S2. Cases selected had a clinical and pathological diagnosis of well-differentiated pNET, expressed at least one of the three neuroendocrine immunohistochemical protein markers (chromogranin A, synaptophysin or CD56) and were surgically resectable at initial diagnosis. Genomic DNA was analyzed from 47 tumors using deep hybridization capture sequencing of 637 genes (578 genes previously associated with cancer, plus an additional 59 genes with published or predicted significance for NETs; Supplementary Table S3). Only a small number of somatic Single Nucleotide Variants (SNVs) and Indels with putative functional significance were identified (Fig. 1; Supplementary Table S5). However, further analysis of this sequence data identified substantial CN gains and losses, in some cases associated with LoH of large regions of the tumor genome (Fig. 2). RNA expression was analyzed from 55 tumors using Affymetrix microarrays, informing the interpretation of these somatic mutations and CN DNA changes.

**Figure 1.**
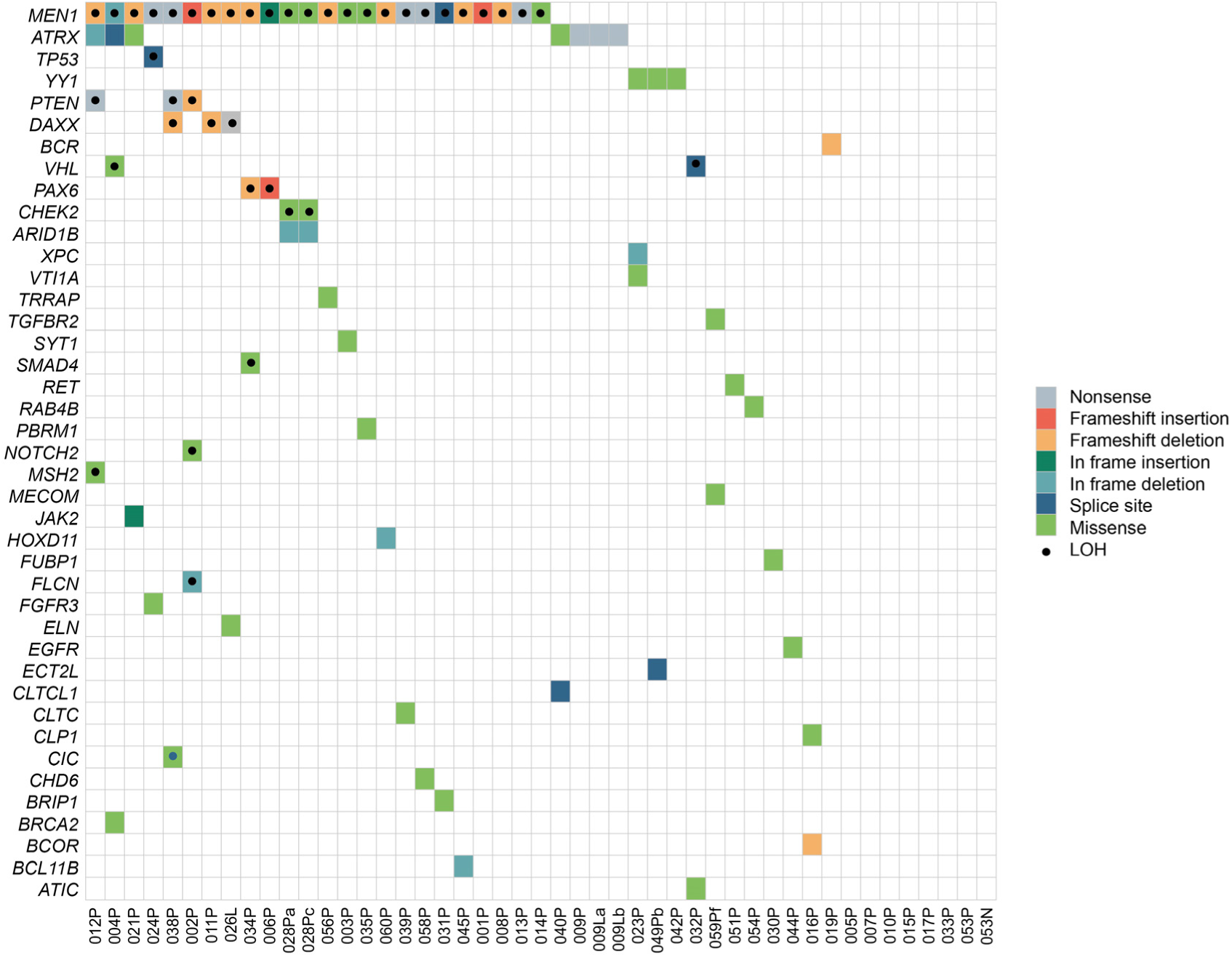
The mutational landscape of pNETs. Coding region somatic non-synonymous SNVs/indels, large deletions and intronic mutations within 2bp of splice sites with any putative functional significance (see Methods) are shown. Tumors are indicated in columns and genes in rows. Colored squares indicate mutation type with dots indicating that loss of the remaining wild type allele (LoH) could be confirmed for the locus through changes in both allele frequency of germline heterozygous SNPs and normalized relative regional sequence depth in tumor vs. normal samples. Multiple tumors from the same patient are indicated by letters after the patient-specific tumor codes (suffix P, N, or L indicates primary pancreas, lymph node or liver metastasis, respectively). In some tumors, there were no detectable mutations in the 637 genes covered by the targeted sequencing panel.

**Figure 2.**
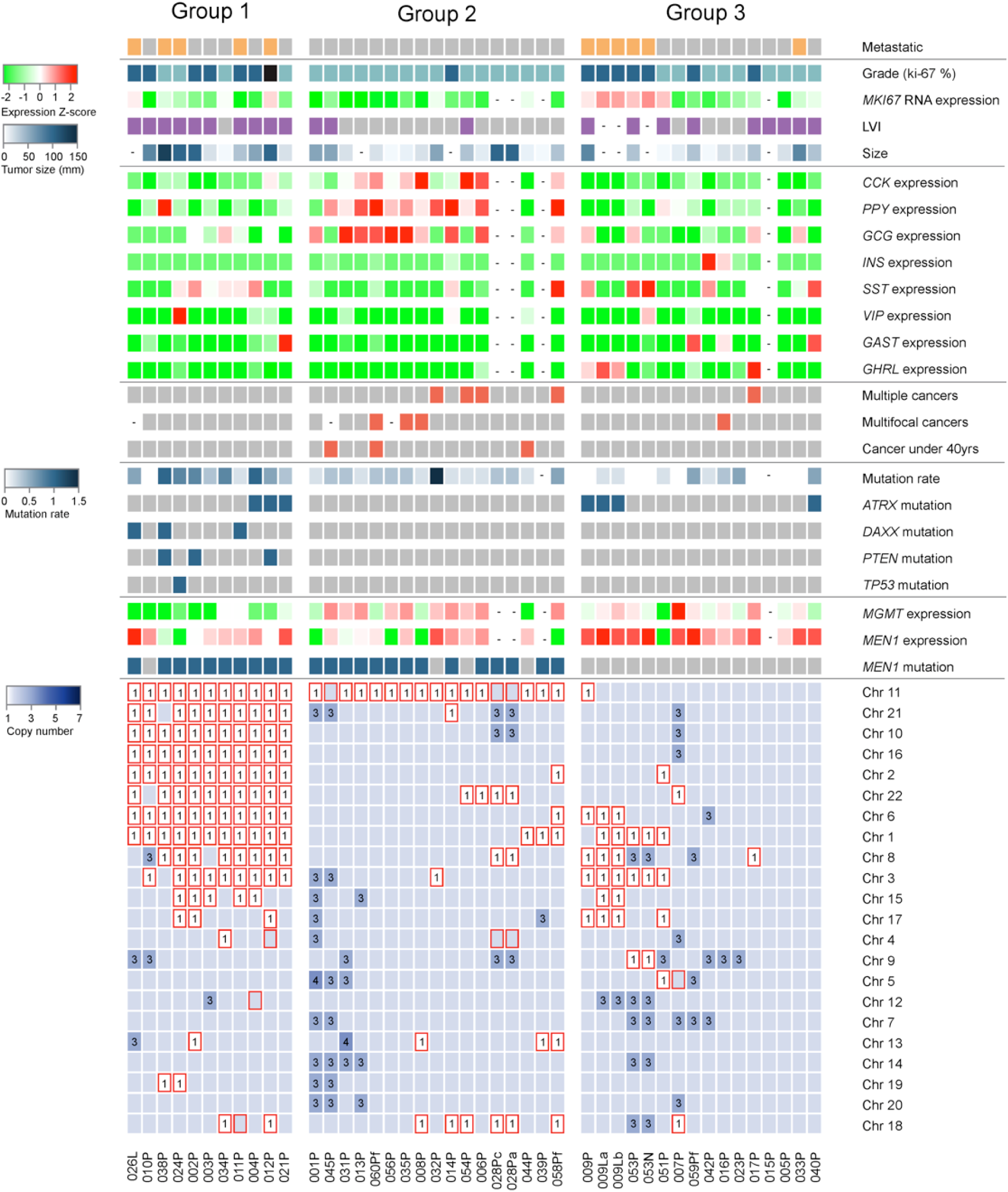
The genomic landscape of pNETs is dominated by aneuploidy. Tumors are shown in columns and genomic and pathological features in rows. Row 1: metastatic tumors are shown in orange. Row 2: grade 1 (Ki-67 ≤2%) is shown in light blue, grade 2 (Ki-67 2-20%) in dark blue and grade 3 (Ki-67 >20%) in black. Row 3: *MKI67* RNA expression Z-score across tumors (green-red color key to left). Dashes indicate that no expression data was available for specific tumors. Row 4 shows the histological identification of lymphovascular invasion (LVI) in purple, tumors without LVI are colored grey. Row 5 shows tumor size (diameter in cm) on a white-blue scale (white-blue color key to left). Rows 6-13 shows expression Z-scores across tumors of the following RNAs (green-red color key to left of row 3): *CCK*, *PPY*, *GCG, INS, SST, VIP, GAST* and *GHRL*. Rows 14-16 show: multiple cancers of any type in the same individual, multifocal pNETs and pNETs arising at under 40 years of age, respectively indicated by red boxes. Row 17 shows the number of functionally significant exonic mutations on a white-blue scale (white-blue color key to left). In rows 18-21 blue squares indicate somatic mutations in the four listed genes. Rows 22 and 23 show the expression of *MGMT* and *MEN1* mRNA (Z-scores, green-red color key to left of row 3). Row 24: Somatic mutations in *MEN1* are shown in blue. In the large bottom panel, coloring of blocks indicates the dominant inferred CN for each autosome in each tumor based on combined information from: ADTEx analysis, relative somatic read counts at germline heterozygous positions and normalized read counts in 3kb tiles across the genome. LoH (irrespective of CN) is indicated by red boxes. Unmarked blue boxes indicate an inferred chromosome CN of 2 and numerals indicate CN when CN ≠ 2.

Additional total RNA and mRNA sequence analysis, methylation microarray analysis and low coverage whole genome sequencing (WGS) was undertaken for the first 12 tumors in this series (summarized in Supplementary Table S4). The WGS confirmed the CN changes revealed by the targeted sequencing. In addition, non-negative Matrix Factorization (NMF) mutational signature analysis of the aggregated WGS data from these 12 pNETs revealed a novel G:C>T:A signature (Supplementary Fig. S1A and B) similar to that recently described in pNETs by Scarpa et al (2).

### Aneuploidy defines the molecular landscape of pNETs and alters gene expression

Primary pNETs are frequently aneuploid (Fig. 2), with 80% (33/41) having LoH of ≥ 1 chromosome and 27% (11/41) having LoH of ≥ 8 chromosomes (Fig. 3). In the aneuploid pNETs, whole chromosome CN was associated with whole chromosome mean RNA expression, shown for 12 pNETs (001P-012P) that had been analyzed by both expression microarray (Fig. 4A and B) and RNAseq (Fig. 4C and D). Although one pNET (009P) appeared to have a low negative association between whole chromosome CN and whole chromosome mean RNA expression, this was due to segmental intra-chromosomal CN variation (Fig. 4E), explaining the low correlation observed at whole chromosome level.

**Figure 3.**
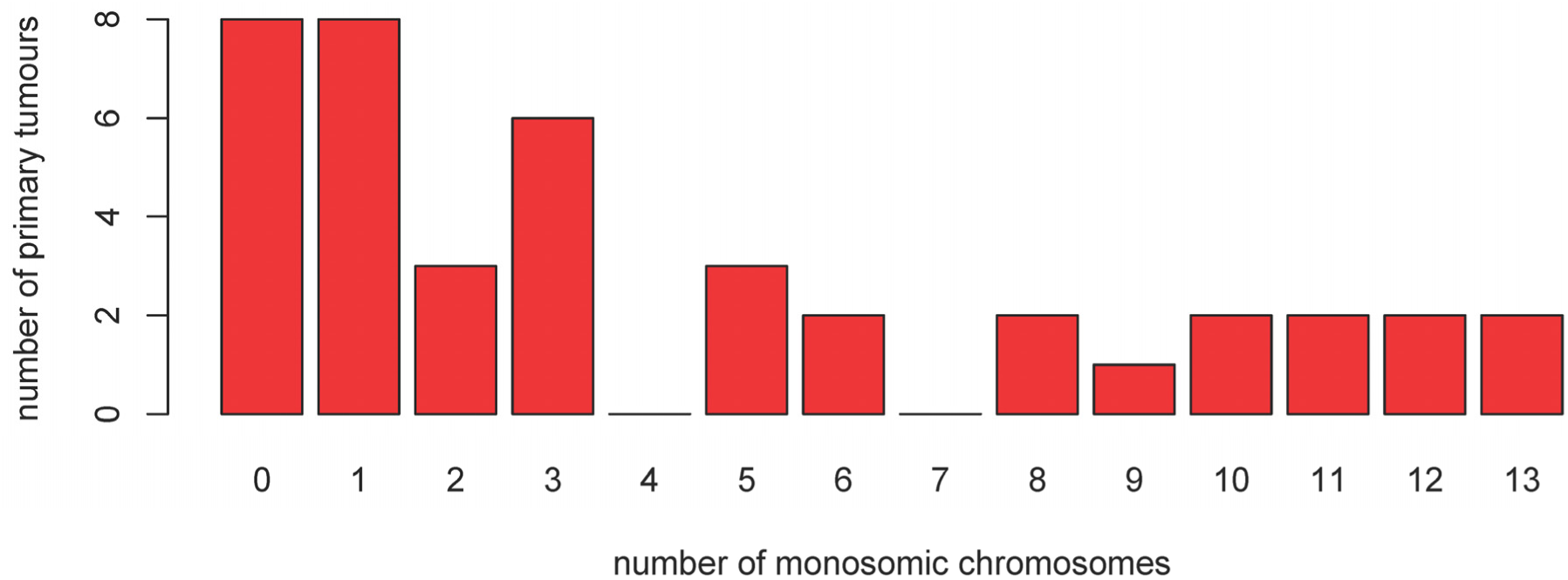
pNET aneuploidy is extensive but varies between tumors. Histogram shows the number of monosomic chromosomes (i.e. both whole chromosome LoH and CN=1) in individual primary tumors.

**Figure 4.**
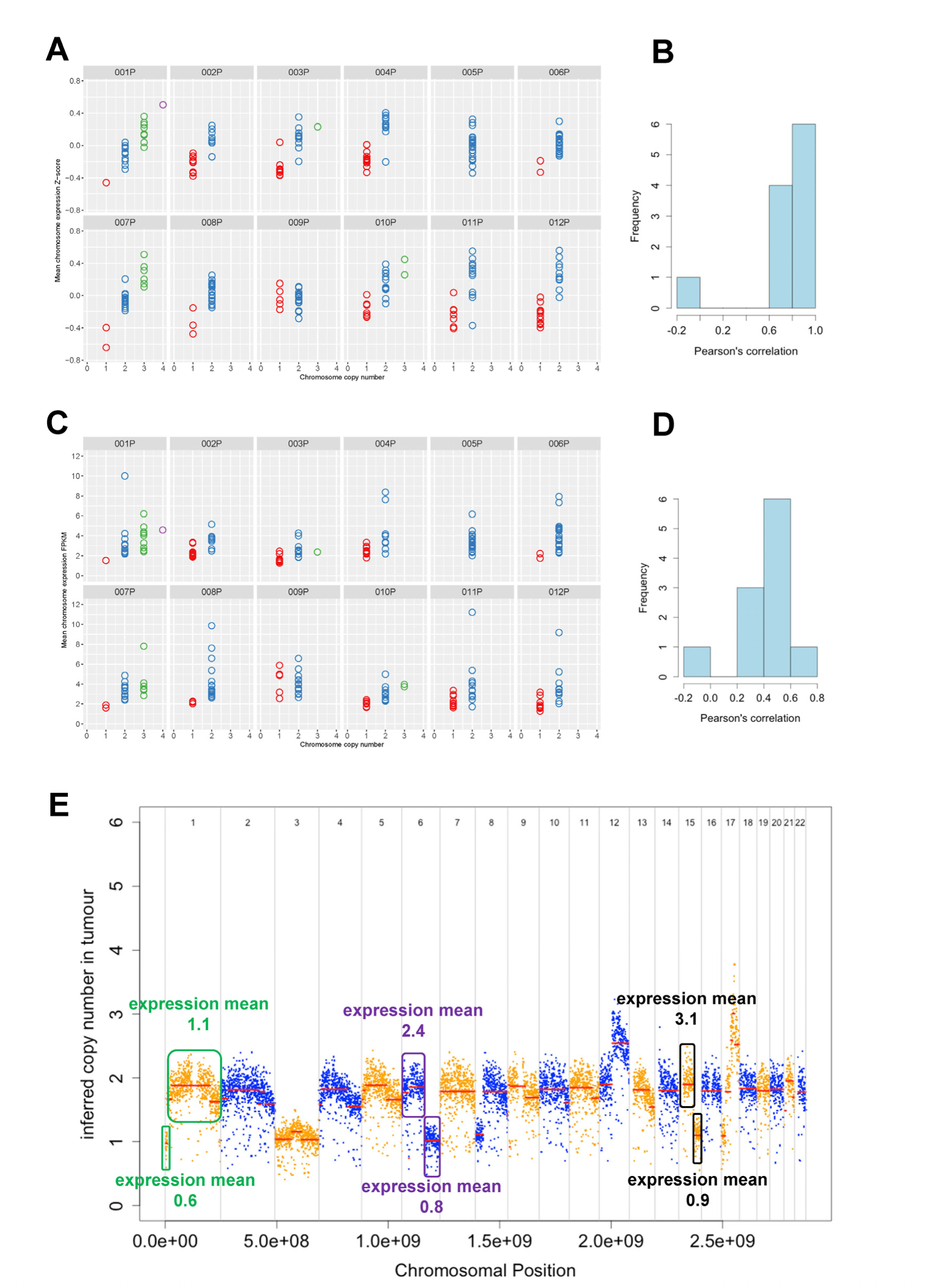
Correlation between DNA copy number and RNA expression for tumors 001P - 012P. Graphs compare whole chromosomal CN (x-axis) to mean chromosomal RNA expression based on **(A)** microarray data or **(C)** RNAseq data (y-axis). Each panel represents a different tumor and each circle represents a different chromosome in that tumor. Histogram of Pearson correlation between CN and **(B)** microarray RNA expression or **(D)** RNAseq RNA expression in each tumor. **(E)** CN across the genome of the tumor 009P that had negative CN-expression correlation (seen at left of histograms in panels B and D). This intra-chromosomal analysis confirms the association between CN and RNA expression seen at whole chromosome level in the other pNETs. Chromosomal segments with specific CN aberrations are shown in colored boxes, with mean RNA expression within each of these segments based on Fragments Per Kilobase of transcript per Million mapped reads (FPKM) shown.

### Patterns of stereotyped aneuploidy and gene expression are associated with clinical behavior

Three distinct pNET groups based on CN change emerged (annotated above Fig. 2). In 11 pNETs (labelled Group 1 in Fig. 2) there was a recurrent pattern of LoH affecting the same 10 chromosomes (1, 2, 3, 6, 8, 10, 11, 16, 21, 22), which has been previously described (2, 6). This idiosyncratic pattern of aneuploidy occurred in the context of somatic *MEN1* mutation in 10 of 11 tumors, an *ATRX* or *DAXX* variant was present in six, with additional *PTEN*, *MSH2* and *TP53* mutations present in five. RNA expression analysis showed that *MGMT* (encoding DNA repair protein O-6-Methylguanine-DNA Methyltransferase) was generally expressed at lower levels in these 11 tumors than in other tumors (t-test *P* = 0.01). Microarray methylation analysis showed *MGMT* gene promoter methylation was relatively consistent across pNETs with no significant correlation to expression (data not shown). Therefore, differential *MGMT* gene methylation, which has been described in other tumor types (12), is unlikely to be the dominant mechanism causing lower *MGMT* RNA expression in this group of pNETs. Instead, one copy of chromosome 10 (the location of *MGMT*) was lost in all 11 tumors in this cluster, suggesting haploinsufficiency as a more likely mechanism for reduced *MGMT* expression (Fig. 2).

Tumors in Group 1 had generally less favorable outcomes; five of the 11 tumors in this group had metastasized, this group contained the only three patients who progressed during the study, and all but one tumor had lymphovascular invasion (LVI) on pathological examination. In contrast, tumors in Group 2 (Fig. 2) were characterized by *MEN1* mutation and chromosome 11 LoH but no recurrent LoH of 10 chromosomes. This group had relatively favorable pathological and clinical outcomes; all had low expression of proliferation-associated RNAs, all but one of the 17 tumors in this group were low grade (Ki-67 <2%), only three had LVI and none metastasized. Several patients in this group had a clinical history loosely associated with inherited cancer predisposition (multiple cancers, multifocal pNETs, and age ≤40yrs). Nine of the 13 tumors in this group for which RNA expression data was available had detectable *GCG* (glucagon) expression. Group 3 (Fig. 2) was characterized by a lack of *MEN1* gene mutation, contained tumors with variable patterns of aneuploidy (ranging from none to extensive) and variable clinical outcomes.

### pNETs have few somatic driver mutations or structural genomic lesions

pNETs have very few detectable somatic variants (Fig. 1, Supplementary Table S5) compared to other tumor types (13) and only one pNET in this study had more than one variant per MB of exons detected (Fig. 5). However, bi-allelic *MEN1* inactivation was common; 81% of tumors with somatic chromosome 11 LoH had a putative pathogenic variant in the remaining *MEN1* allele (Fig. 2). These variants were distributed across the entire *MEN1* coding region (Supplementary Fig. S1C). Analysis of *MEN1* expression showed that nonsense and frameshift variants were associated with reduced *MEN1* RNA abundance (Supplementary Fig. S1D). This suggests that processes such as nonsense-mediated decay may contribute to reduced abundance of MENIN protein in *MEN1* mutant tumors, in addition to the pathogenic changes introduced by these mutations altering MENIN protein structure and function.

**Figure 5.**
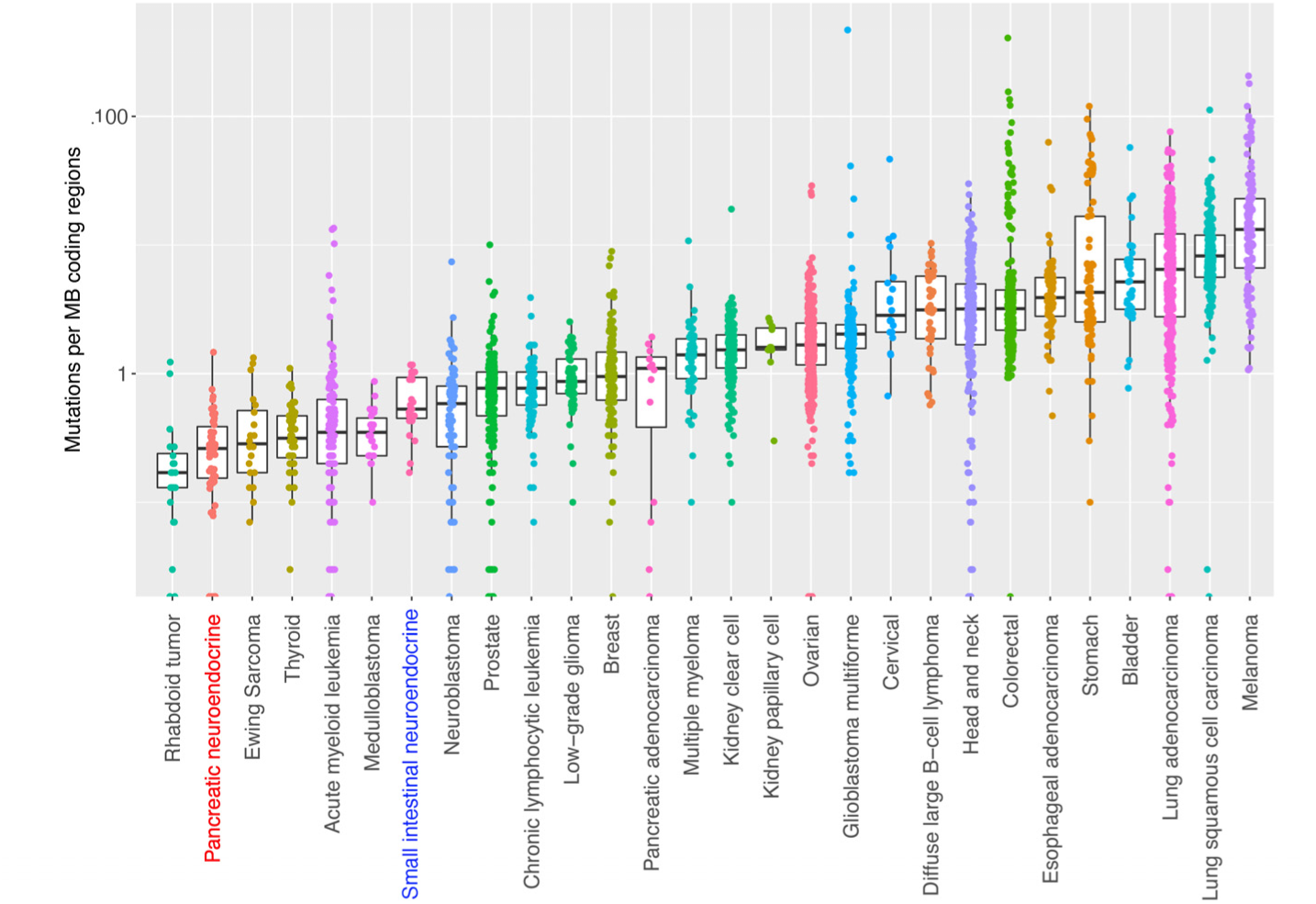
Number of mutations in pNETs. pNETs have relatively low somatic mutation frequency compared to other tumor types; box plots show the coding region mutation rate of pNETs compared to the coding region mutation rates described by Lawrence et al. (13) in other tumors analyzed by WES or WGS.

Methylation analysis showed no clear correlation between *MEN1* gene methylation and RNA expression in 15 tumors tested, suggesting that methylation is not an important regulator of *MEN1* expression in pNETs.

MutSig analysis (13) identified *MEN1* as the only statistically significant cancer driver gene across this pNET cohort, although previously described variants in a small number of other tumor suppressor genes were seen in multiple tumors (Fig. 1 and 2) including *ATRX*, *DAXX*, *PTEN, YY1* and *VHL*. Private variants in 34 other genes were detected in single patients, some of which were clinically interesting as previously described predictive biomarkers for specific therapies, including bi-allelic inactivation of *MSH2* and mono-allelic variants in genes such as *RET, JAK2, FGFR3* and *BRCA2* (Fig. 1). Although these variants were classified as functionally significant using combinations of variant effect databases (see Methods) and many are considered clinically actionable using current assessment tools (Supplementary Fig. S2), genomic analyses suggested that most were passenger mutations rather than drivers of tumorigenesis. For example, examination of variants in genes encoding tyrosine kinases that have matching small molecule inhibitors found no corroborating *JAK2* (patient 021), *FGFR3* (patient 024) or *RET* (patient 051) gene expression/pathway changes. However, the mutation pathogenicity of variants in one tumor was corroborated by the RNA expression data – tumor 002P had a somatic frameshift mutation in *PTEN* (a phosphoinositide 3-kinase (PI3K)/AKT/mTOR signaling inhibitor) as well as a non-frameshift deletion and LoH in the *FLCN* gene (encodes the mTOR complex 2 inhibitor folliculin). Microarray analysis of the expression of RNAs downstream of PI3K suggested significant activation of PI3K/AKT/mTOR pathways in this tumor, consistent with the mutations (Supplementary Fig. S3).

### Pseudohypoxia determines the expression profiles of some pNETs

Tumors from six patients (8 samples) had high expression of a subset of the hypoxia-activated RNAs. These tumors also tend to have more rapid proliferation based on both *MKI67* RNA expression (Supplementary Fig. S4A) and immunostaining (7 of these tumors were grade 2 NETs with Ki-67 immunostaining in 3-20% of nuclei). However, further genomic analysis showed that two of the eight tumors had somatic *VHL* variants with LoH, and tumors from other patients had high *VHL* gene methylation associated with significantly low *VHL* RNA expression (Supplementary Fig. S4B). This suggests that in the majority of the pNETs analyzed, tumor hypoxia gene expression profiles are due to pseudohypoxia caused by disrupted VHL function rather than true hypoxia.

### Germline variants may become significant in the context of extensive aneuploidy

We were able to exclude with high confidence any functionally relevant germline variants in the following genes previously associated with NETs: *MEN1, RET, TSC1, TCS2, PTEN, NF1, CDKN1B, IPMK, MAX, NF1*, *NTRK1*, *SDHA*, *SDHB*, *SDHC*, *SDHD, MUTYH* and *VHL*. However, there were 173 germline variants in 66 genes not traditionally associated with NETs that were predicted to disrupt protein function (Supplementary Table S6). The list of genes affected was significantly enriched for genes associated with DNA repair (GO:0006281, *P* = 6×10^−9^) using the Panther web tool (14). Eight of these variants appeared to become unopposed when their remaining normal allele was lost by somatic LoH (Supplementary Table S7). These 8 variants had ~1:1 ALT:REF allele ratios in germline DNA, and all had somatic LoH in tumor DNA and corresponding tumor ALT:REF allele ratios of ≥ 1.5. In these tumors, the degree of loss of the remaining normal allele was consistent with the proportion of tumor comprising somatic cells rather than stroma. As an example, tumor 014P had a chr11:108098576_C/G variant in *ATM* with an ALT:REF allele ratio of 0.9 in the germline but 2.8 in the tumor due to LoH (Fig. 6A). This variant has a population frequency of 0.007 in the ExAc database (15) and leads to a p.Ser49Phe amino acid substitution. Although Clinvar indicated this was a variant of uncertain significance, analysis using IPA and GeneSetDB indicated that numerous RNAs with expression dependent on ATM function were downregulated in this tumor (Fig. 6B), consistent with somatic LoH exposing a pathogenic germline variant causing somatic loss of ATM activity.

**Figure. 6.**
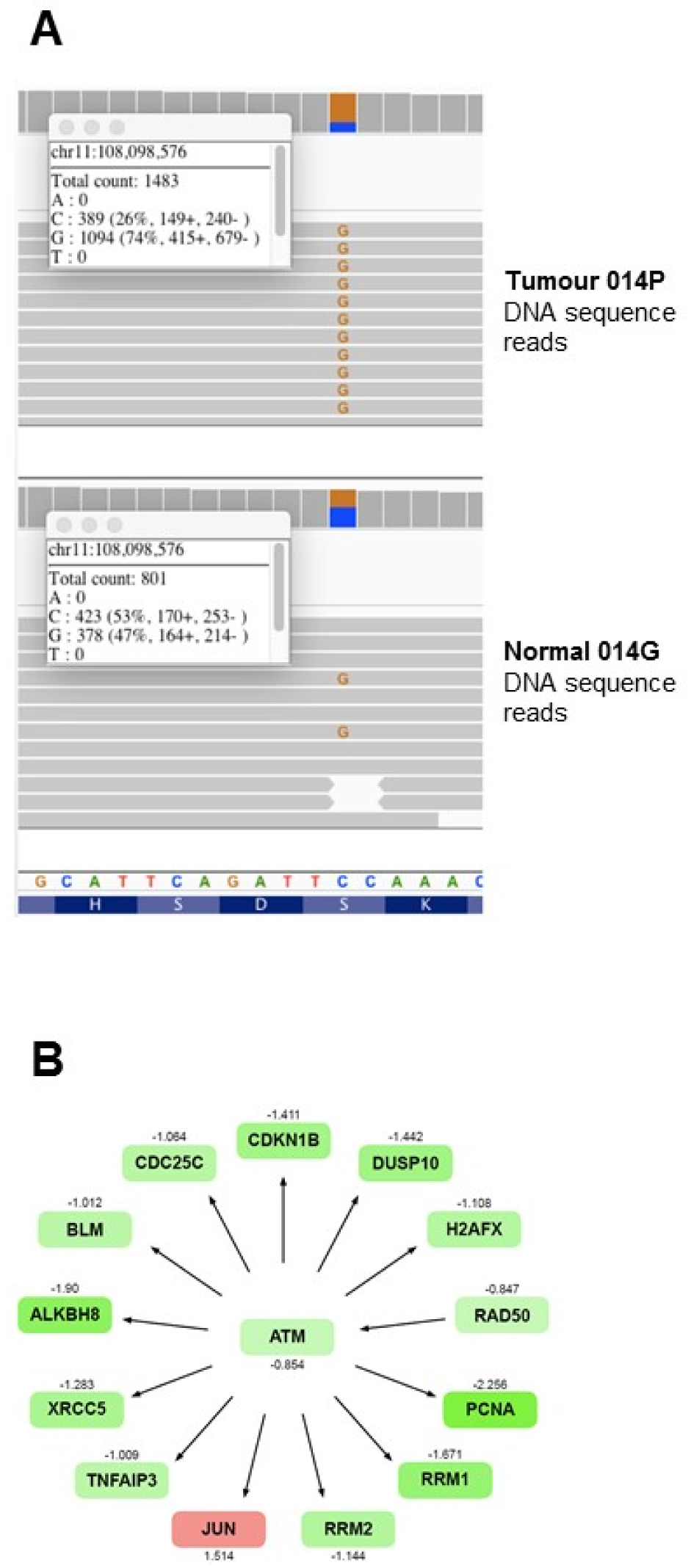
Somatic LoH may expose germline heterozygous variants. **(A)** Example of a germline heterozygous mutation in the *ATM* locus that becomes unopposed in the tumor 014P due to somatic LoH. **(B)** IPA analysis indicates that in this tumor expression of numerous RNAs that are normally up-regulated by activity of the ATM complex is generally reduced, suggesting reduced ATM function. Shades of red and green indicate the degree of up-and down-regulation of RNAs, respectively, with Z-score expression values shown below each node.

### Immune, proliferative and hormone expression characteristics of pNETs

The pVAC-Seq neoantigen prediction framework (16) putatively identified only one tumor (012P) with a mutation capable of generating a potential neoantigen consistent with the patient's HLA haplotype. This low incidence of predicted neoantigens is not surprising, given the low somatic mutation rate in pNETs (Fig. 5). There was no association between grade and somatic variant frequency, suggesting that tumor grade is not determined by single gene events in pNETs.

Pancreatic endocrine hormone RNA expression analysis was conducted to characterize the expression of sub-clinical functioning (e.g., insulinoma, glucagonoma) and "non-functioning" pNETs. Approximately two thirds of pNETs had detectable expression of the RNA encoding at least one hormone, despite the absence of symptoms reported clinically by patients. Expression of the RNAs *INS* (encodes Insulin) and *IAPP* (encodes Amylin) appeared correlated, in line with their known co-production by pancreatic islet β cells (Supplementary Fig. S5A). The expression of a set of RNAs not usually noted to be co-expressed in the same islet cells (*GCG*, *PPY* and *CCK*) also appear correlated. Tumors expressing *VIP* (encodes Vasoactive Intestinal Peptide) RNA did so exclusively (Supplementary Fig. S5A).

Tumors expressing *GHRL* (encodes ghrelin, produced by ε; cells) RNA also did so exclusively, and methylation analysis found that high *GHRL* expression (in the three tumors from patient 009) had low mean methylation of CpG islands in the *GHRL* gene promoter, suggesting that dysregulated methylation may have contributed to *GHRL* expression in this patient (Supplementary Fig. S5B). Differential gene promoter methylation was not associated with RNA expression for any pancreatic endocrine hormones other than ghrelin in the 15 tumors assessed. *INS* RNA expression was only associated with a clinical diagnosis of ‘insulinoma’ (biochemically proven hypoglycemia caused by pNET insulin secretion) in a subset of tumors, and in some tumors there appeared to be *INS* RNA expression without a documented clinical syndrome (Supplementary Fig. S5A).

## Discussion

### Unusual genomic lesions

By combining multiple types of genomic analysis we have shown that pNETs develop through a range of unusual oncogenic mechanisms. Although more than half of pNETs have biallelic loss of *MEN1*, the overall frequency of somatic SNVs, Indels and structural DNA variants was low, with small numbers of tumors carrying tumor suppressor variants in *ATRX*, *DAXX*, *VHL*, *PTEN*, *YY1* and *PAX6*. These variants generally concord with previously published analyses of mutations in pNETs (1, 2, 4, 17, 18).

Instead of mutation, it appears that most pNETs are defined by variable and extensive aneuploidy. For example, approximately 80% of pNETs in this series had lost a copy of ≥1 chromosome and a recurrent pattern of aneuploidy was observed in some pNETs, which carried LoH of an identical set of 10 chromosomes (Fig. 2), therefore affecting thousands of genes. Somatic haploinsufficiency is a plausible mechanism by which this LoH may contribute to pNET development, supported by the striking association we demonstrate between RNA expression and CN (Fig. 4). Given the large number of genes affected on these 10 chromosomes (≥9,500), it is difficult to identify gene sets or pathways significantly enriched above what could occur by chance. Nevertheless, a range of tumor suppressor genes are now thought to drive tumor development through haploinsufficiency rather than by simple mutation (19) and aneuploidy can disrupt entire signaling pathways (20), especially those that depend on precise stoichiometry of protein subunits (21). It is possible that development of some pNETs may be driven predominantly by aneuploidy, analogous to chromosome 5q-deleted myelodysplastic syndrome in which haploinsufficiency without specific mutation appears to drive the neoplasia (22). Understanding the origin, selection in tumor populations and clinical consequences of recurrent aneuploidy, such as we see here, remains a key challenge for modern cancer biology.

Haploinsufficiency is a tenable direct cause for the low *MGMT* RNA expression in pNETs with the recurrent pattern of 10-chromosome loss, since heterozygous *MGMT* +/-mouse tissues have significantly reduced O-6-Methylguanine-DNA Methyltransferase activity (23). Since low *MGMT* function is one of the determinants of response to alkylating agents such as temozolomide (24), these pNETs with low *MGMT* expression may potentially respond to temozolomide therapy, and this aneuploid genotype may have contributed to variations in response to temozolomide in published series (25, 26). We also show an example where somatic LoH renders a pathogenic heterozygous germline variant in *ATM* unopposed, thereby inactivating its downstream tumor suppressor pathways (Fig. 6). Careful inspection of the germline in each patient found no evidence of traditional syndromic NET-associated mutations, or the recently recognized *MUTYH* germline mutations (2) in our cohort.

Distinct patterns of pancreatic islet hormone expression were seen in the majority of pNETs and may indicate the cell of origin of these tumors. Although RNA expression might not translate into protein expression, high expression of RNAs encoding hormones in most pNETs analysed suggests that clinicians should be aware of under-diagnosis of subtle secretory syndromes. A recent study of 26 insulinomas found mutations, copy number changes and focal allelic imbalances in genes significantly enriched for epigenetic regulators (11). However, we could not replicate these observations in the 12 clinically-defined insulinomas in our pNET cohort. Neither the mutations we detected in these 12 pNETs, nor the genes affected by aneuploidy, showed significant enrichment for epigenetic factors listed in EpiFactors database (27). Interestingly, we also observed one patient with three metastatic tumors with high *GHRL* (encodes ghrelin) (28) RNA expression and promoter hypomethylation.

### Distinct genomic landscapes, putative oncogenic mechanisms and clinical features of two pNET subsets

By combining CN, somatic variant analysis and expression analysis we hypothesize that there are distinct oncogenic mechanisms driving two subsets of pNETs (Fig. 2), summarized in Fig. 7. The first subset, Group 1, are pNETs with *MEN1* mutation coupled with recurrent loss of 10 chromosomes, the cause of which remains unclear. This subset generally had unfavorable grade 2 and 3 histology, all but one patient had LVI and five of the 11 tumors in this group metastasized, and *MGMT* loss through apparent haploinsufficiency may favor the use of temozolomide. The second subset, Group 2, contained pNETs with *MEN1* mutation and chromosome 11 LoH but few other changes in chromosomal CN - none of this group never went on to metastasize and all but one had favorable low grade histology (Ki-67 <2%). In addition, all of this second subset had low expression of proliferation-associated RNAs, only three of the 17 tumors in this subset had LVI and most expressed the RNA encoding glucagon. In this subset, the decision to leave tumors un-resected could be considered in the setting of a clinical trial, thus avoiding the complications and long-term morbidity of surgery for these patients.

**Figure 7:**
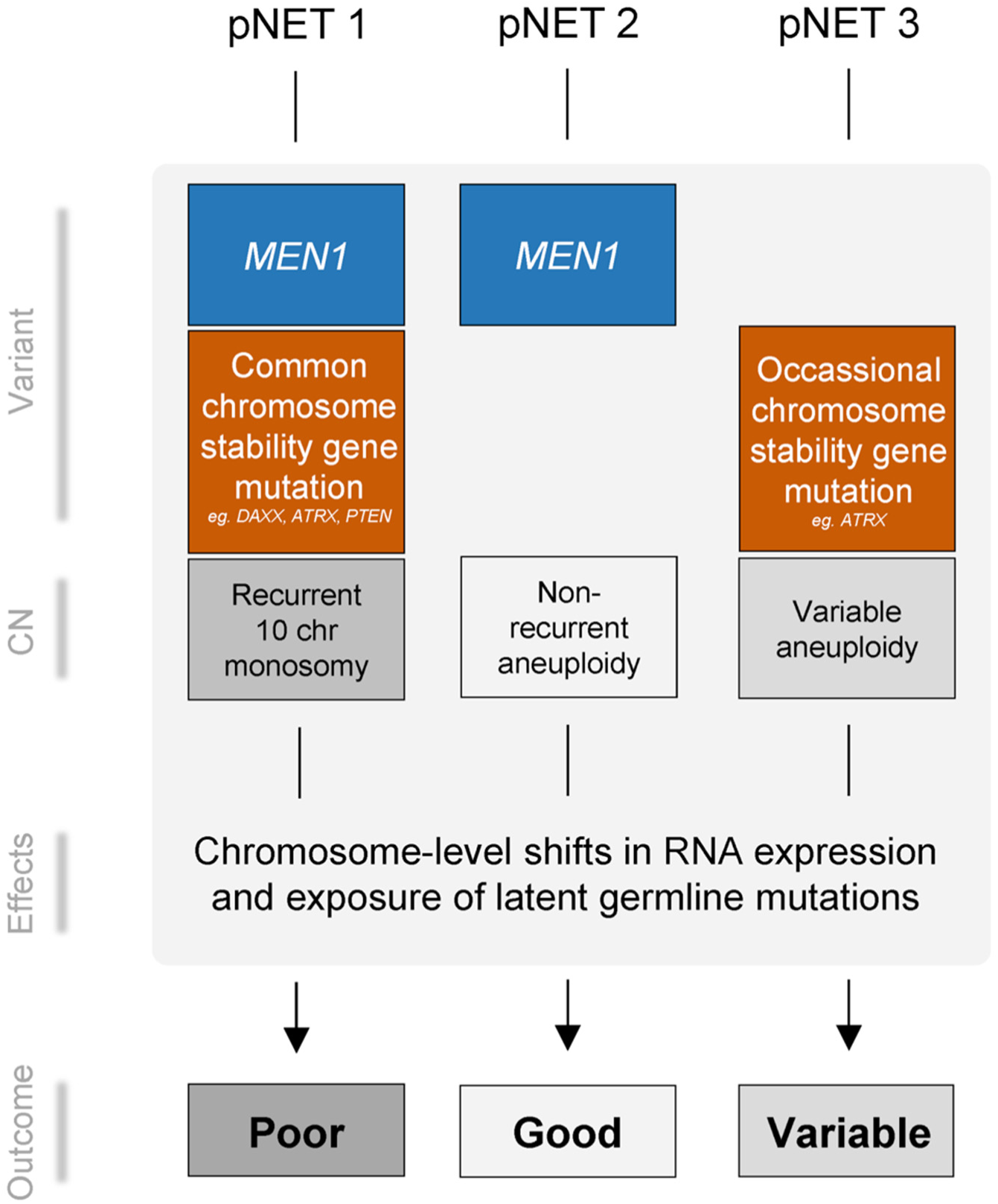
Integrated genomic, pathological and clinical categorization of pNETs. Genomic features within pNETs identified three groups: Group 1 generally have *MEN1* mutation and chromosome 11 loss, sporadic mutation of genes associated with chromosomal instability, recurrent loss of ten specific chromosomes leading to extensive disruption of gene expression, and reduced *MGMT* expression. These genomic features are strongly associated with high tumor grade and size, LVI and more frequent metastases. Group 2 have *MEN1* mutation and chromosome 11 loss but no recurrent loss of ten chromosomes. They have universally low tumor grade, size and LVI, many express *GCG* RNA and importantly, this group have no metastases. Group 3 are characterized by no *MEN1* mutation, with variable aneuploidy, clinical and pathological features.

### Conclusion

Aneuploidy appears to be a fundamental event driving pNET tumorigenesis, possibly by altering gene expression on a global scale and exposing pathogenic germline variants, leading to signaling pathway dysregulation. Tumors with few mutations and massive aneuploidy, such as many pNETs, will not be amenable to simple precision oncology paradigms that match drugs to single gene mutations; these will require more nuanced approaches that take account of oncogenic drivers that affect more than just individual genes.

## Methods

### Tumor sample collection and processing

Surgically resected, fresh frozen and FFPE specimens were collected from the Cancer Society Tissue Bank, University of Otago, NZ and Auckland Region Hospitals under New Zealand Health and Disability Ethics committee approvals 13/NTA/69 and 13/NTB/173 (for further information on sample handling, see supplementary methods). Matched blood and normal adjacent tissue, where possible >20mm distant, were used as germline controls for the fresh frozen and FFPE cases respectively. Frozen tissues were processed to isolate genomic DNA (Macheray Nagel; Nucleospin Tissue kit; #740952) and total RNA (Ambion miRvana RNA isolation kit; Thermo Fisher Scientific; AM1560). FFPE tissues were macro-dissected on slides to maximize tumor cellularity and gDNA and RNA isolated by QIAamp DNA FFPE kit (Qiagen; #56404) or Ambion RecoverAll kit (Thermo Fisher Scientific; AM1975). Whole blood/buffy coat and dried FTA blood spots (6 × 3mm punches) were extracted using QIAamp DNA mini kits (#51102, #51304). All isolation kits were used as per the manufacturer’s instructions. Nucleic acid quality and quantity were determined by Agilent Tapestation and Qubit Fluorometry respectively.

### RNA analysis

All microarray hybridization and sequencing machine runs for RNAseq were performed as a service by New Zealand Genomics Ltd. For RNAseq, 100ng RNA from tumors 001P-012P were used as templates to prepare separate total and mRNA libraries using TruSeq Stranded Total RNA Sample Prep kit with Ribo-Zero gold and TruSeq Stranded mRNA Sample Prep kits, respectively. They were sequenced as a multiplex of 6 samples per HiSeq lane using V3 chemistry, 2×100PE reads. The total RNA and mRNA sequencing reads were trimmed using cutadapt (29) v1.9.1 to remove leftover adapters, any reads with Phred score of <30, and read pairs where either read was <50bp after trimming. Reads were aligned using Bowtie 2 (30) with recommended settings. Aligned total RNA reads and mRNA reads were merged using Picard MergeSamFiles before gene and transcripts abundance quantified using RSEM (31). Fusion genes were searched for using TopHat-Fusion (32).

For microarray expression analysis, Affymetrix PrimeView Human Gene Expression arrays were used (perfect-match-only microarrays with ~530,000 probes covering ~36,000 transcripts). 100ng RNA was labelled using the Affymetrix SensationPlus FFPE method according to manufacturer instructions, before hybridization to the gene chips, washing and scanning. QC was performed using Affymetrix Expression Console and in-house R (33) scripts to visualize probe signal distributions relative to control signals. Data for one tumor (015P) was discarded due to very low tumor-derived signals relative to spiked-in control signals, and remaining tumor data was quantile normalized in R using RMA (34), as implemented in the R ‘affy’ package (35). To remove any systematic FFPE-vs-fresh frozen sample biases, for each probe the mean signal within all FFPE tumors was subtracted from each individual FFPE tumor’s signal, with an identical adjustment performed within the fresh frozen tumor group. A comparison between the results of RNAseq and microarray analysis revealed relatively high concordance and is shown in Supplementary Fig. S6.

For all visualizations, expression values for each probe set were transformed into Z-scores (by mean centering the data then expressing variation above and below the mean on a scale of standard deviation) and all analysis of Probe set differential expression used the LIMMA R package (36). Differentially expressed probe sets were tested for enrichment of particular functional categories or pathways using IPA (37), GATHER (38) and GeneSetDB (39). Stromal content was estimated from sequence data using ADTEx (40), with Immune subtype abundance in the tumors estimated using the Cibersort (41) and Estimate (42) methods. mRNA expression for specific gene sets was visualized combined with clinical and pathological information in a web dashboard using the R Shiny package, using a modification of the heatmap.2 function of the gplots package (43) to draw annotated heatmaps.

### Methylation analysis

500ng of gDNA from each of the 001P-012P, 009La, 009Lb tumors was bisulfite converted as per manufacturer instructions for the EZ DNA methylation kit (Zymo; D5001). Samples were labelled and hybridized as a service by AgResearch Ltd, GenomeNZ section, New Zealand onto Illumina Infinium Human 450k methylation arrays. Data was visualized and QC performed using the ChAMP package (44), which also provided estimates of tumor gene CN. All methylation BeadChips passed QC standards recommended in the ChAMP documentation. Methylation β values were subsequently extracted from the idat files with the RnBeads (45) package in R using the hg19 human genome assembly and mean aggregation of each of: promoters, CpG islands and genes. Measurements were filtered using the Greedycut algorithm; background was subtracted using the noob method of the methylumi package before signal intensity normalization using the SWAN method of the minfi package (46). mRNA expression data (microarray) and gene methylation data were linked through Ensemble gene ID of the respective platform annotation files. A local MySQL database was generated and queried through statistical filters (significance of correlation and level of expression) to identify significantly anti-correlated/correlated methylation and mRNA expression, using the RSQLite package.

### DNA sequencing and data analysis

WGS libraries were generated for tumors 001P-012P using a Rubicon ThruPLEX-FD kit with 3-50ng of input DNA. For shallow WGS each WGS library was run 1 sample per lane of HiSeq (but split over multiple lanes) with V3 Chemistry 2×100PE reads. WES enrichment was performed using the Agilent SureSelect V5 + UTR systems on the above libraries and run in a multiplex of 3 per lane as per the WGS analysis. For targeted sequencing, the NimbleGen SeqCap EZ comprehensive cancer panel was used (Roche; #4000007080 − a ~4Mb design that targets 578 cancer related genes). An additional custom SeqCap EZ choice panel (Roche NimbleGen 06266282001) covering 59 additional genes with a capture space of 354Kb was also designed (Custom NET panel) (Supplementary Table S3). This was completed according to manufacturer instructions, and as further described in the supplementary methods.

Sequencing reads were quality trimmed using cutadapt v1.9.1 to remove left over Illumina specific adapters and any reads with Phred score of <30. Read pairs were removed if either read had an after trimming length of <50bp. Reads were aligned to UCSC hg19 reference genome using BWA-mem (47) with default settings and duplicated reads then removed using Picard v2.1.0 MarkDuplicates (http://broadinstitute.github.io/picard/). Aligned reads with minimum mapping quality of 1 were selected using Samtools (48). Due to the high sequence depth achieved by target capture, maximum depths were set to 9,000, the Samtools per-Base Alignment Quality calculation was removed and tumor purity was set to 50%. Finally, the strand filter was removed as it is not applicable to target-captured data. The R SomaticSignatures package (49) was used to identify and plot mutational signatures using non-negative matrix factorization (Supplementary Fig. S1), with the R SomaticCancerAlterations package (50) used to retrieve TCGA tumor somatic variant data to provide comparable mutational signatures from non-NET tumor cohorts. Putative origins of mutational signatures are based on information in Alexandrov et al. (51).

### Variant calling and annotation

Somatic SNVs and Indels were primarily detected using the VarScan2 (52) v2.3.7 somatic workflow. Somatic variants were detected in parallel using Strelka (53) and qSNP (54) without filtering with default settings – all SNVs and Indels described in this paper could be detected using all three methods. Neither Varscan's germline nor somatic p-value filters were used. VCF files were extensively annotated using PERL scripts modified from ANNOVAR (using ANNOVARs ljb26 database) with additional annotations from The Cancer Genome Interpreter (https://www.cancergenomeinterpreter.org). Variants detected in presence of supplementary reads were additionally annotated with a custom flag in the original VCF file using vcf-annotate in VCFtools (55). Somatic variants were then filtered in real time, while visualizing the effects of the filtering, using a web-based variant visualization dashboard based on the R VariantAnnotation and Shiny (56) packages. This dashboard was also used to generate many of the figures in this paper. It used ggplot2 (57) for graphics, the httr package to provide real time links to the IGV genome browser (58), and functions modified from the R DNAcopy package (59) for circular binary segmentation subroutines and for visualization of segmental CN aberrations and corresponding segmental B-allele frequency changes. Post-calling filtering used the following criteria: Normal tissue and tumor read depth at the site of the mutations ≥ 50, ≥ 10 tumor sequences showing the mutation, the site of mutation is not within the Encode Dac Mapability black list, and ≤ 2 reads corresponding to the mutation were found in the germline sample. All somatic variants that passed these filters were visually validated in IGV. Germline SNVs and Indels were detected using the VarScan2 v2.3.7 germline workflow. Mutation plots and lollipop plots in Figs. 1, 2 and Supplementary Fig. 1C were generated using modifications of the waterfall and lolliplot functions, respectively, of the GenVisR R package (60).

### Coding region mutation rate analysis

The pNET coding region mutation rate was compared to the rates described by Lawrence *et al*. (13) in other tumor types, which had been analyzed by either WGS or WES. Mutation frequencies were calculated in each pNET based on the numbers of coding region mutations found in regions of the genome with ≥ 50x sequence coverage. Note that pNET mutation frequencies may be overestimated in this analysis since, compared to the WGS or WES analysis used to calculate coding region mutation rates for the other tumor types, the hybridization capture analysis used here is enriched for cancer-associated genes, which may be more likely to carry mutations.

### Copy-number and structural variant analysis

CN analysis was performed using the deep targeted sequencing data for all tumors and separately using the WGS data available for tumors 001P-012P. CN variation was first visualized by counting the number of reads mapped to 3 kb tiles of the hg19 human genome using bedtools multicov, then analyzed using the DNACopy R package. For each 3kb tile, all raw counts were log2-transformed, normalized using loess splining and log ratios between tumor and normal were calculated, the log ratios were smoothed, segmented (circular binary segmentation) and visualized. B allele frequencies were also analyzed across the tumor genomes to combine with CN information in order to identify combination of intra-tumoral heterogeneity and unbalanced chromosomal amplification. To do this required identification of germline heterozygous positions, which was based on: 0.4 < proportion of ALT reads in germline <0.6 <u>and</u> probability ≥0.95 of the observed germline sequence reads being sampled from a population of reads where ALT and REF alleles were equally common, calculated using a binomial distribution. Somatic CN aberrations were also analyzed in parallel using the Varscan2 CN pipeline, supplemented by statistical analyses using ADTEx (40), Titan (61) and for some tumors CN information from Infinium Methylation BeadChips were analyzed using ChAMP. WGS, WES and SeqCap aligned BAM files for patients 001 – 012 were merged using Picard MergeSamFiles before somatic structural variants were analyzed using MANTA (62), Delly2 (63) and GRIDSS (64) using default settings; in the other tumors, structural variants were analyzed using these three packages from SeqCap data alone.

### Data Materials and Availability

All data will be made available via European Genome-phenome Archive (https://www.ebi.ac.uk/ega/home). Data access will be granted via a ‘data access committee’ who release data to researchers who apply and meet specific guidelines.

## Acknowledgements

We thank all patients that donated samples and clinical data to our study. We also thank Stefan Bohlander (University of Auckland) for his help with genomic data analysis and interpretation, Ruellyn Cockcroft, Simon Fu, Rebecca Jones and Ole Schmeidel (Auckland DHB), Dean Harris (Canterbury DHB), Richard Issacs (Mid-Central DHB and Jeremy Rossaak (Bay of Plenty DHB) for clinical input, Siobhan Conroy and Avril Hull of the Unicorn Foundation for coordinating patient input and support, Melissa Firth for input to the Register set up, Maui Hudson and Helen Wihongi for cultural advice and David Jansen for help with translation of key aspects of our project into Te Reo Maori, Malcolm Legget for ongoing advice as Chair of the Translational Medicine Trust, Ramon Gallego, Aaron Jeffs, Purvi Kakadiya, Rebecca Laurie, John Markham, Les McNoe, John Pearson, Thalanie Purdy, Puja Sharma, Bruce Tsai, Nic Waddell, and Liam Williams for help in the design and interpretation of laboratory studies. This work was financially supported by the Translational Medicine Trust.

## Author contributions

BL was the co-principal investigator on the project and clinical lead, CB coordinated all laboratory work and handling of clinical samples, KP led the development of the register of patient information, coordinated the identification and procurement of clinical samples, PT conducted many of the bioinformatics analyses, supported by VF and NP, MB provided bioinformatics input and expert advice, SF, PS, TR and SJ conducted DNA and RNA extraction and laboratory analysis, MLY and NK conducted complete pathology review of all samples and, along with RR and MY helped with sample identification and procurement, PY provided clinical genetic review of our analysis, EC and BW worked on the development of the patient register, BR provided clinical annotation of patient samples and clinical input, KH and PR provided cultural advice and support and helped in the development of a governance framework for the project to support the rights of indigenous persons, JK, PJ, RC, SC, HM, JW, AM, RB and AB provided help with sample procurement and clinical review of the manuscript, ME, CJ and DD provided clinical expertise in the interpretation of our analyses, NK provided bioinformatics and statistical input, SG provided expert review of the genomic analysis and advice re project design and further work, MF was co-principal investigator on the project and provided clinical input, CP was co-principal investigator on the project and provided genomics and bioinformatics expertise.

## Competing interests

None.

## References

1. Jiao Y, et al. (2011) DAXX/ATRX, MEN1, and mTOR pathway genes are frequently altered in pancreatic neuroendocrine tumors. Science 331(6021):1199–1203.

2. Scarpa A, et al. (2017) Whole-genome landscape of pancreatic neuroendocrine tumours. Nature.

3. Cao Y, et al. (2013) Whole exome sequencing of insulinoma reveals recurrent T372R mutations in YY1. Nat Commun 4:2810.

4. Heaphy CM, et al. (2011) Altered telomeres in tumors with ATRX and DAXX mutations. Science 333(6041):425.

5. Marinoni I, et al. (2014) Loss of DAXX and ATRX are associated with chromosome instability and reduced survival of patients with pancreatic neuroendocrine tumors. Gastroenterology 146(2):453–460 e455.

6. Nagano Y, et al. (2007) Allelic alterations in pancreatic endocrine tumors identified by genome-wide single nucleotide polymorphism analysis. Endocr Relat Cancer 14(2):483–492.

7. Missiaglia E, et al. (2009) Pancreatic endocrine tumors: expression profiling evidences a role for AKT-mTOR pathway. Journal of Clinical Oncology 28(2):245–255.

8. Larsson C, Skogseid B, Oberg K, Nakamura Y, & Nordenskjold M (1988) Multiple endocrine neoplasia type 1 gene maps to chromosome 11 and is lost in insulinoma. Nature 332(6159):85–87.

9. Stefanoli M, et al. (2014) Prognostic relevance of aberrant DNA methylation in g1 and g2 pancreatic neuroendocrine tumors. Neuroendocrinology 100(1):26–34.

10. Sadanandam A, et al. (2015) A Cross-Species Analysis in Pancreatic Neuroendocrine Tumors Reveals Molecular Subtypes with Distinctive Clinical, Metastatic, Developmental, and Metabolic Characteristics. Cancer Discov 5(12):1296–1313.

11. Wang H, et al. (2017) Insights into beta cell regeneration for diabetes via integration of molecular landscapes in human insulinomas. Nature Communications 8.

12. Gerson SL (2004) MGMT: its role in cancer aetiology and cancer therapeutics. Nat Rev Cancer 4(4):296–307.

13. Lawrence MS, et al. (2013) Mutational heterogeneity in cancer and the search for new cancer-associated genes. Nature 499(7457):214–218.

14. Mi H, Muruganujan A, & Thomas PD (2013) PANTHER in 2013: modeling the evolution of gene function, and other gene attributes, in the context of phylogenetic trees. Nucleic Acids Res 41(Database issue):D377–386.

15. Lek M, et al. (2016) Analysis of protein-coding genetic variation in 60,706 humans. bioRxiv.

16. Hundal J, et al. (2016) pVAC-Seq: A genome-guided in silico approach to identifying tumor neoantigens. Genome Med 8(1):11.

17. Bosman FT, Carneiro F, Hruban RH, & Theise ND (2010) WHO classification of tumours of the digestive system (World Health Organization).

18. Corbo V, et al. (2010) MEN1 in pancreatic endocrine tumors: analysis of gene and protein status in 169 sporadic neoplasms reveals alterations in the vast majority of cases. Endocrine-related cancer 17(3):771–783.

19. Davoli T, et al. (2013) Cumulative haploinsufficiency and triplosensitivity drive aneuploidy patterns and shape the cancer genome. Cell 155(4):948–962.

20. Durrbaum M & Storchova Z (2016) Effects of aneuploidy on gene expression: implications for cancer. FEBS J 283(5):791–802.

21. Santaguida S & Amon A (2015) Short-and long-term effects of chromosome mis-segregation and aneuploidy. Nat Rev Mol Cell Biol 16(8):473–485.

22. Komrokji RS, Padron E, Ebert BL, & List AF (2013) Deletion 5q MDS: molecular and therapeutic implications. Best Pract Res Clin Haematol 26(4):365–375.

23. Glassner BJ, et al. (1999) DNA repair methyltransferase (Mgmt) knockout mice are sensitive to the lethal effects of chemotherapeutic alkylating agents. Mutagenesis 14(3):339–347.

24. Thon N, Kreth S, & Kreth FW (2013) Personalized treatment strategies in glioblastoma: MGMT promoter methylation status. Onco Targets Ther 6:1363–1372.

25. Fine RL, et al. (2013) Capecitabine and temozolomide (CAPTEM) for metastatic, well-differentiated neuroendocrine cancers: The Pancreas Center at Columbia University experience. Cancer Chemother Pharmacol 71(3):663–670.

26. Strosberg JR, et al. (2011) First-line chemotherapy with capecitabine and temozolomide in patients with metastatic pancreatic endocrine carcinomas. Cancer 117(2):268–275.

27. Medvedeva YA, et al. (2015) EpiFactors: a comprehensive database of human epigenetic factors and complexes. Database 2015:bav067.

28. Corbetta S, et al. (2003) Circulating ghrelin levels in patients with pancreatic and gastrointestinal neuroendocrine tumors: identification of one pancreatic ghrelinoma. The Journal of Clinical Endocrinology & Metabolism 88(7):3117–3120.

29. Martin M (2011) Cutadapt removes adapter sequences from high-throughput sequencing reads. EMBnet.journal 17:10.

30. Langmead B & Salzberg SL (2012) Fast gapped-read alignment with Bowtie 2. Nat Methods 9(4):357–359.

31. Li B & Dewey CN (2011) RSEM: accurate transcript quantification from RNA-Seq data with or without a reference genome. BMC Bioinformatics 12:323.

32. Kim D & Salzberg SL (2011) TopHat-Fusion: an algorithm for discovery of novel fusion transcripts. Genome Biol 12(8):R72.

33. Team R (2014) R: A language and environment for statistical computing. R Foundation for Statistical Computing, Vienna, Austria. 2013.

34. Irizarry RA, et al. (2003) Summaries of Affymetrix GeneChip probe level data. Nucleic Acids Res 31(4):e15.

35. Gautier L, Cope L, Bolstad BM, & Irizarry RA (2004) affy--analysis of Affymetrix GeneChip data at the probe level. Bioinformatics 20(3):307–315.

36. Smyth GK (2004) Linear models and empirical bayes methods for assessing differential expression in microarray experiments. Stat Appl Genet Mol Biol 3:Article3.

37. Kramer A, Green J, Pollard J Jr., & Tugendreich S (2014) Causal analysis approaches in Ingenuity Pathway Analysis. Bioinformatics 30(4):523–530.

38. Chang JT & Nevins JR (2006) GATHER: a systems approach to interpreting genomic signatures. Bioinformatics 22(23):2926–2933.

39. Araki H, Knapp C, Tsai P, & Print C (2012) GeneSetDB: A comprehensive meta-database, statistical and visualisation framework for gene set analysis. FEBS Open Bio 2:76–82.

40. Amarasinghe KC, et al. (2014) Inferring copy number and genotype in tumour exome data. BMC Genomics 15:732.

41. Newman AM, et al. (2015) Robust enumeration of cell subsets from tissue expression profiles. Nat Methods 12(5):453–457.

42. Yoshihara K, et al. (2013) Inferring tumour purity and stromal and immune cell admixture from expression data. Nat Commun 4:2612.

43. Warnes M & Bolker B (2016) Package 'gplots'. Various R Programming.

44. Morris TJ, et al. (2014) ChAMP: 450k Chip Analysis Methylation Pipeline. Bioinformatics 30(3):428–430.

45. Assenov Y, et al. (2014) Comprehensive analysis of DNA methylation data with RnBeads. Nat Methods 11(11):1138–1140.

46. Aryee MJ, et al. (2014) Minfi: a flexible and comprehensive Bioconductor package for the analysis of Infinium DNA methylation microarrays. Bioinformatics 30(10):1363–1369.

47. Li H (2013) Aligning sequence reads, clone sequences and assembly contigs with BWA-MEM. arXiv preprint arXiv 00:3.

48. Li H, et al. (2009) The Sequence Alignment/Map format and SAMtools. Bioinformatics 25(16):2078–2079.

49. Gehring JS, Fischer B, Lawrence M, & Huber W (2015) SomaticSignatures: inferring mutational signatures from single-nucleotide variants. Bioinformatics 31(22):3673–3675.

50. J G (2016) SomaticCancerAlterations: Somatic Cancer Alterations. R package version 1.10.0.

51. Alexandrov LB, et al. (2013) Signatures of mutational processes in human cancer. Nature 500(7463):415–421.

52. Koboldt DC, et al. (2012) VarScan 2: somatic mutation and copy number alteration discovery in cancer by exome sequencing. Genome Res 22(3):568–576.

53. Saunders CT, et al. (2012) Strelka: accurate somatic small-variant calling from sequenced tumor-normal sample pairs. Bioinformatics 28(14):1811–1817.

54. Kassahn KS, et al. (2013) Somatic point mutation calling in low cellularity tumors. PLoS One 8(11):e74380.

55. Danecek P, et al. (2011) The variant call format and VCFtools. Bioinformatics 27(15):2156–2158.

56. Winston Chang JC, JJ Allaire, Yihui Xie, Jonathan McPherson (2016) shiny: Web Application Framework for R. R package version 0.14.2.

57. Wickham H (2009) ggplot2: elegant graphics for data analysis. Springer New York.

58. Thorvaldsdottir H, Robinson JT, & Mesirov JP (2013) Integrative Genomics Viewer (IGV): high-performance genomics data visualization and exploration. Brief Bioinform 14(2):178–192.

59. Olshen VESaA (2016) DNAcopy: DNA copy number data analysis. R package version 1.48.0.

60. Skidmore ZL, et al. (2016) GenVisR: Genomic Visualizations in R. Bioinformatics 32(19):3012–3014.

61. Ha G, et al. (2014) TITAN: inference of copy number architectures in clonal cell populations from tumor whole-genome sequence data. Genome Res 24(11):1881–1893.

62. Chen X, et al. (2016) Manta: rapid detection of structural variants and indels for germline and cancer sequencing applications. Bioinformatics 32(8):1220–1222.

63. Rausch T, et al. (2012) DELLY: structural variant discovery by integrated paired-end and split-read analysis. Bioinformatics 28(18):i333–i339.

64. Cameron DL, et al. (2017) GRIDSS: sensitive and specific genomic rearrangement detection using positional de Bruijn graph assembly. bioRxiv:110387.

